# Electro-diffusion modulation of synaptic input in dendritic spines using deconvolved voltage sensor time series

**DOI:** 10.1101/097279

**Authors:** J. Cartailler, T. Kwon, R. Yuste, D. Holcman

## Abstract

The inward current flowing inside the post-synaptic terminal of a neuron modulates transiently the membrane voltage potential. Most of the excitatory connections are made on dendritic spines characterized by a large variability in their geometry.How the voltage in a spine is modulated by geometry remains elusive due in part to the absence of direct measurements. To understand the spine voltage-current relation, we develop here a model for the voltage and we use it to extract electrical properties from live cell imaging data. We first deconvolve the genetically encoded voltage sensor expressed in hippocampal neurons and then use electro-diffusion theory, to compute the electric field and the ionic flows induced in a dendritic spine.The ionic flow is driven by the electric field coupled to the charge densities that interact through the non-cylindrical spine geometry. We determine the I-V relation and conclude that the spine effective resistance is mostly determined by the neck geometry. This modulation of synaptic inputs by the spine neck is significantly larger than what was expected from traditional cable models. Thus modulating the postsynaptic current can be achieved by changing the number of receptors or by altering the spine geometry which independently affects the transformation of current into voltage.

**Significance statement:** Dendritic spines are geometrical structures receiving most of the excitatory transmission, yet how they modulate voltage from the synaptic current is not clear due to their submicron small size and specific non-cylindrical geometry. We study here the conversion of the synaptic current into voltage modulated by the spine geometry. Our approach is based on the electro-diffusion theory and we show that the spine neck is the main resistance filter, while the voltage is maintained constant in the head. Finally, we extract the effective resistance using a deconvolution of the genetically encoded voltage indicators expressed in hippocampal neurons. The present approach allows studying the electrical properties of many other structures such as glial small protrusions, cilia, and many others.

## Introduction

Neurons communicate through synaptic microdomains, where an input current generates a voltage change in the post-synaptic neuron. This voltage change reflects the strength of the synaptic connection between two interacting neurons [1, 2, 3, 4, 5] and depends on two components: the first is the number of glutamatergic receptors for excitatory neurons and the second is the geometry of the post-synaptic terminal. However the relative contribution between these two factors is still unclear. For example, the postsynaptic structure is often a dendritic spine, the geometry of which can modulate the time scale of diffusion [6, 7, 8, 9]. In parallel, increasing the number of receptors on that terminal leads to a larger synaptic current. But, given the influence of electric fields on ionic species, diffusion alone is not sufficient to interpret the synaptic response driven by electro-diffusion, involving the coupling of diffusion to the voltage gradient. Electrodiffusion was applied successfully for studying ionic fluxes and gating of voltage-channels [10, 11], because at the nanometer scale, cylindrical symmetry of a channel model reduces to a one-dimension for the electric field and charge densities in the channel pore [12, 13]. Moreover, the current in the synaptic cleft has already been shown to reflect the coupling between moving ions and the local electrical field [14, 15].

In a dendritic spine, voltage changes during the synaptic response should be generated by the interactions between the ionic flow and the spine geometry. Dendritic spines are heterogenous microdomains at the limit of optical resolution and for that reason, voltage changes were estimated for many years using modeling and numerical simulations of the cable equations, the basis for the Hodgkin-Huxley model [16]. This approach is however not appropriate for spines because the micro-geometry is significantly different from that of a cable. Also, cable theories break down when applied to small neuronal compartments, such as dendritic spines, because they assume spatial and ionic homogeneity. Linear approximations of electro-diffusion that couples the electric field with the ionic flow have been used to improve the estimation of the voltage changes in spines, approximated as cylinders of various sizes, but assuming local electroneutrality [4]. Recent advent in monitoring the voltage changes at a sub-micrometer resolution [17, 18, 19] can now reveal the electrical properties of dendritic spines [20, 21], but the heterogeneity of the results [22, 23] and the absence of a robust computational framework and theory to interpret data challenges our understanding of electrical properties of these structures and cellular microdomains in general.

We develop here an electro-diffusion framework to compute the voltage-current relation and the local voltage variations generated by synaptic inputs. We also present a deconvolution procedure to recover the time scale of voltage responses from voltage-sensitive indicator hippocampal neurons. After we present the deconvolution procedure to transform the voltage dye arclight response into voltage dynamics, we use the Poisson-Nernst-Planck theory for electrodiffusion to extract the current flowing in the spine neck and the electrical resistance. Numerical simulations of the voltage drop in the entire spine reveal how a change in the neck length and radius alters the voltage changes in the entire spine. We conclude here that while the numbers and the types of receptors determine the injected current, the geometry of a dendritic spine controls the conversion of current into synaptic voltage.

## Results

### Converting the Arclight signal into a fast voltage response

For these experiments we used optical measurements of Archlight fluorescence to extract voltage estimates of the response of dendritic spines and neighboring dendrites from cultured hippocampal neurons.

The Arclight dye indicator responds to voltage with an intrinsic delay [24], such that a fast voltage response leads to a convolve light response. In that context, a synaptic input entering a dendritic spine generates a light response that needs to be deconvolved in order to recover the electrical time course. As the voltage intensity has already been deconvolved in [24], we focus here on the time dependent response. The basis of the method is to find the kernel of the deconvolution *K*(*t*), which is obtained by comparing the electrophysiological and the light responses in the soma (see Methods). Once the kernel is found, we apply it to recover the voltage dynamics in smaller structure such as spines and portions of dendrites.

Fig. 1A-B show the soma region to be deconvolved. The deconvolution procedure transforms the fluorescence dye (green) signal into the voltage response (black) (Fig. 1B). The deconvolve signal is the dashed line (green) which superimposes with the electrophysiological recordings (continuous black line). We confirm the validity of the method by using the direct convolution (black large arrow) of the electrophysiological recording by the kernel, which leads to a response (black curve) that exactly super-imposes with the fluorescence soma signal (Fig. S1). Regions of interest are shown in Fig. 1C, where we shall deconvolve the fluorescent responses. Indeed, once the kernel of the deconvolution *K*(*t*) is determined from the soma, we shall apply it to deconvolve the signal in smaller structures such as spines, where the signal contains large fluctuations (Fig. 1D-E dashed curves). The results are shown in Figs. 1D-E for the voltage the spine head and dendrite. The procedure to remove the fluctuation is described in the Method section and in Fig. S2. Due to the slow time scale of glutamate uncaging, that can last for hundred of milliseconds, the voltage responses we obtained in the dendritic spines are slower than the expected direct synaptic response.

**Figure 1:**
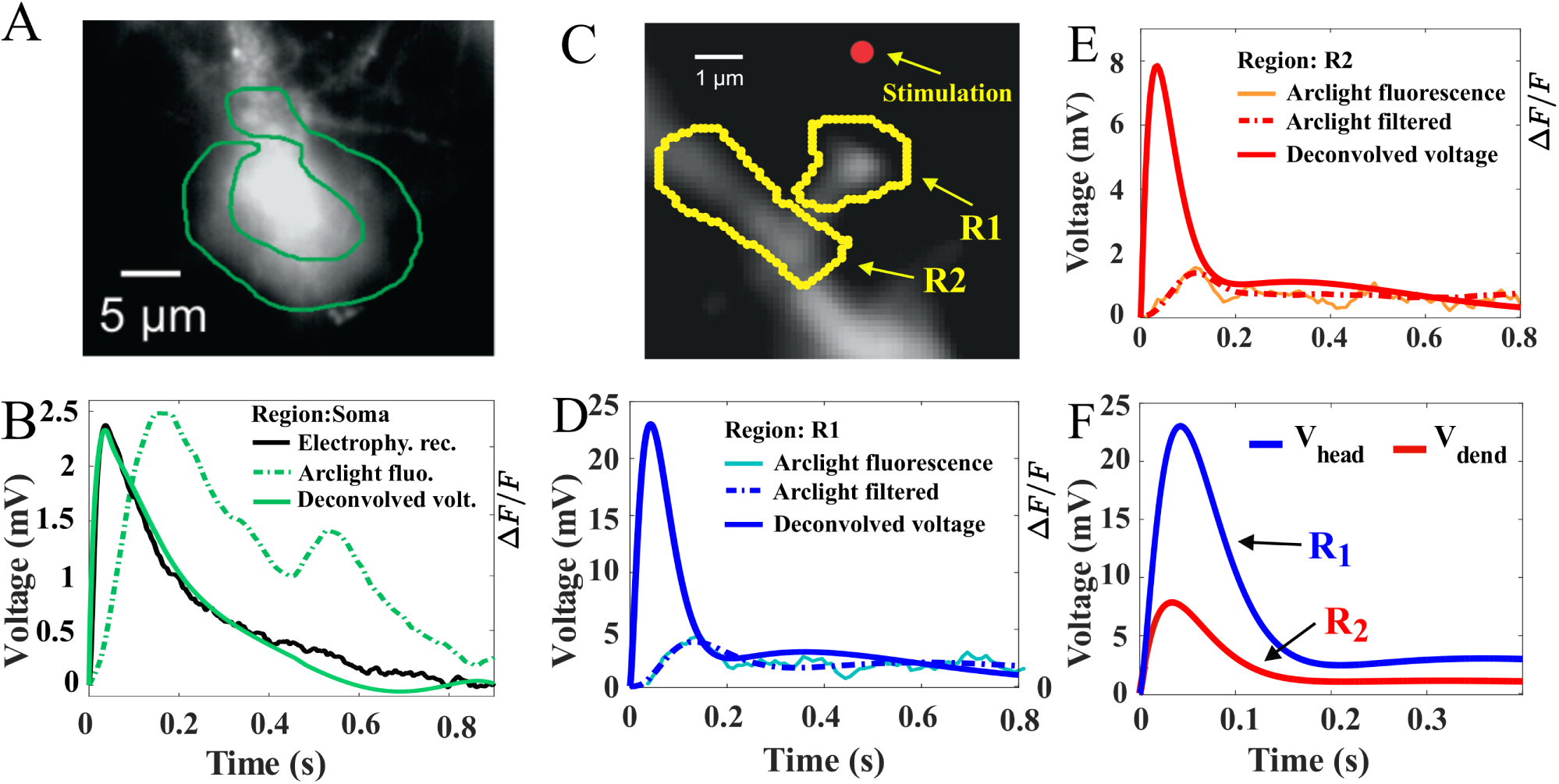
Voltage drop across a dendritic spine measured from Arclight. **A.** A region of interest is selected around the soma to estimate the fluorescence during a synaptic stimulation. **B.** The deconvolution of the fluorescence signal (dashed green) in the soma uses the electrophysiological recording (continuous black) to obtain the deconvolved voltage (continuous green). **C.** Deconvolution of the fluorescence signal in the dendritic spine *R*1 and the dendrite *R*2 using the kernel *K*(*t*) found in the soma deconvolution in (**A-B.**). **D-E.** Deconvolution of the regions of interest (ROIs) *R*1 and *R*2 obtained from a fluorescent Arclight response following a glutamate uncaging protocole (red dot). **F.** Comparison of the filtered and deconvolved voltage signal in the spine head and parent dendrite.

### Electro-diffusion theory for ionic flows in dendritic spines

To interpret the voltage dynamics in a dendritic spine, we use the electro-diffusion model that couples positive *c_p_*(*x*,*t*) and negative *c_m_*(*x*,*t*) charge concentrations with the electrical potential *V*(*x*,*t*). The model is the phenomenological Poisson-Nernst-Planck (PNP) equations, where the ionic flow is driven by diffusion and an electric force. The voltage potential is described by the classical Poisson equation for the charges [25] (see Methods). We use this ensemble of equations to model the flow of ions when a current *I*(*t*) is injected at the entrance of a dendritic spine. While cations can enter or exit the spine domain, we assume that negative charges stay confined. We recall that the electrical potential generated by a flow of ions is defined to an additive constant.

We apply the electrodiffusion approach at the nano-micrometer scale, by reducing the neck geometry to a one dimensional segment (Fig. 2A-B). Due its large size, the dendrite constitutes a ionic reservoir compared to a dendritic spine. We fix in the dendrite the voltage *V* = 0 mV. We thus interpret the potential *V*(*t*) = *V*(*t*,*L*) computed at the end of the neck (Fig. 2A-D) as the difference of potential between the entrance and the exit of the spine neck. To describe the response of an input current *I*(*t*) inside the spine neck, we approximate the thin cylinder as a one dimensional wire of length L (see eqs. 8-9-10).

**Figure 2:**
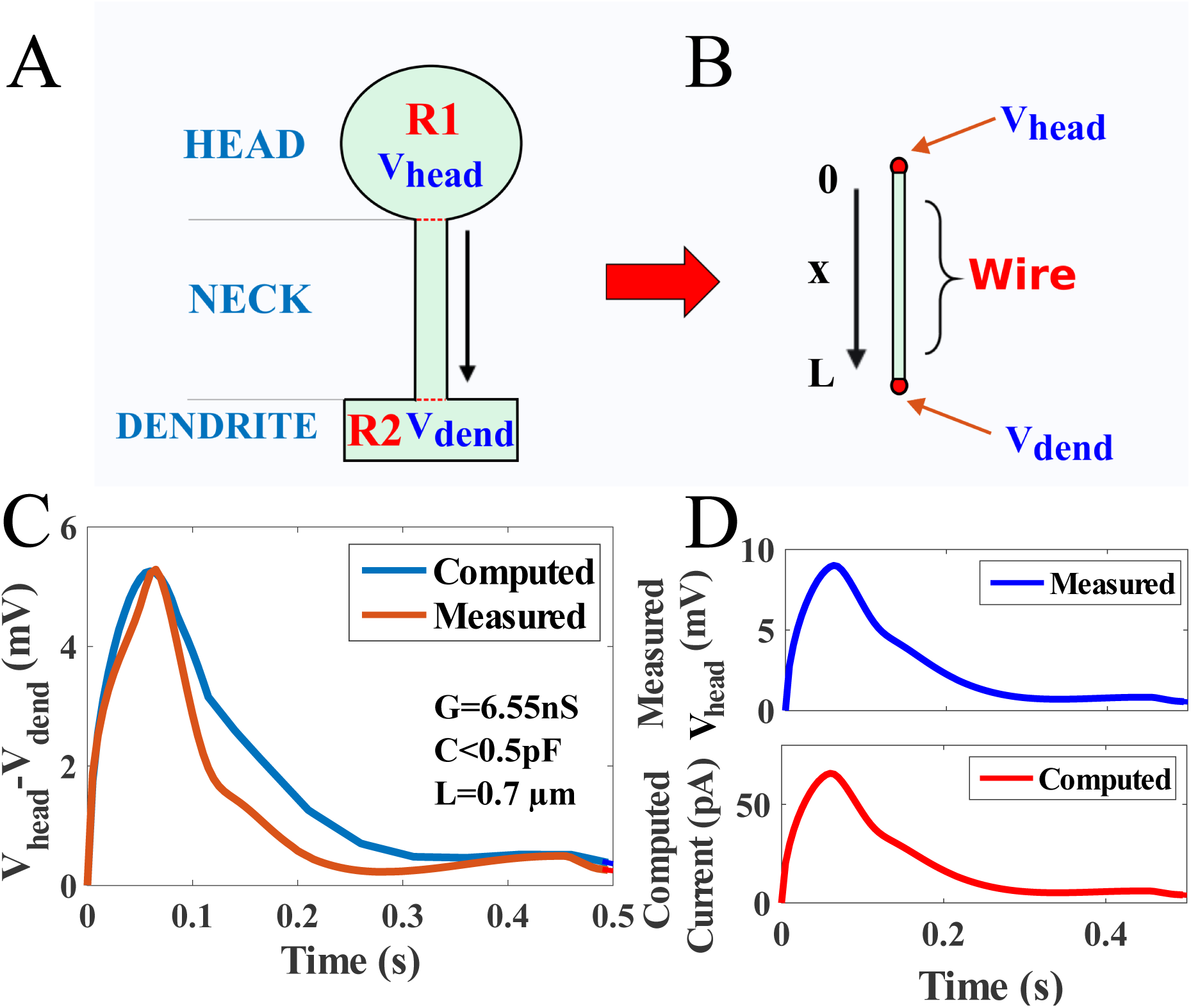
Extracting the electrical parameters of a dendritic spine from a reduced model and voltage dynamics. **A.** Schematic representation of a dendritic spine, divided into three regions: the head *R*1, the neck of length *L* and the close dendrite *R*2. **B.** Reduced geometry of a dendritic spine neck of length *L*, approximated as a dielectric wire. The input is the measured voltage *V*_1_(*t*) of ROIs (of fig. 1) *R*1 at the head (*x* = 0) and we use voltage *V*_2_(*t*) (*R*2) in the parent dendrite (*x* = *L*) as an output for comparison with the numerical computation. **C.** Comparison of measured and computed voltage. **D.** Measured membrane potential (blue) in the spine head is used to compute the ionic current (red) from eq. 1, after the parameters (*C*, *G*) are extracted from the iterative method developed in the Methods

From the classical theory of electricity [25], it is not possible to extract the current passing through a passive devise from the difference of potential when the resistance is unknown. However, by using a model for the current in the spine head, we shall reconstruct the voltage in the neck and recover the current in the entire spine. Indeed, following a synaptic input, the current *I*(*t*) flowing in the spine is driven mostly by positive sodium charges *c_p_*(*x*,*t*). Because there is no direct measurement of this current, we developed here a procedure (see SI) to estimate this current from the measured membrane potential *ϕ*(*t*) in the spine head. We model the current *I*(*t*) as the sum of a resistance and a capacitive:

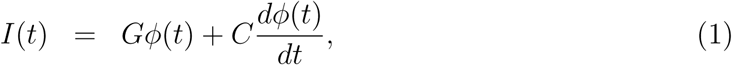

where *G* is the intrinsic conductance of the spine and *C* is the capacitance. To estimate the two constants *C* and *G* and the voltage drop across a spine neck, we solve numerically eqs. 8-9-10 and compare the simulations with the deconvolved Arclight fluorescent voltage response, following glutamate uncaging stimulations at the top of the spine head (Fig. 1C). We solve numerically the PNP equations for the distribution of positive *c_p_*(*x*,*t*) and negative *c_m_*(*x*,*t*) charges, as well as the potential difference *V*(*x*,*t*) (Fig. 2C-D). To estimate the voltage difference Δ *Ṽ*(*t*) across the neck, we grounded the potential to 0 mV at the dendritic shaft (before stimulations, the voltage is described by eq. 9).

To assess whether the potential difference Δ*Ṽ*(*t*) = *V_head_*(*t*) — *V_dend_*(*t*) can be predicted from the electro-diffusion model, we fix the input voltage *ϕ*(*t*) = *V_head_*(*t*). We then compare the voltage obtained by solving eqs. 8-9-10 (Fig. 2C-D) to the measured voltage *V_dend_*(*t*) in region *R*2 (blue) at the dendritic shaft (Fig. 2B). Although region *R*1 includes the head and the neck, we neglected the fluorescence in the neck due to its small thinness ≤ 100*nm*. We found a good agreement between the experimental data and numerical simulations (Fig. 2E) showing that the difference of voltage between the head and dendrite can be predicted from the input voltage *V_head_*(*t*). In addition, we estimated the injected current (Fig. 2D) (see Methods and eq. 1) directly without any direct electrophysiological recordings. We conclude at this stage that the electro-diffusion theory allows estimating the electrical properties (capacitance and resistance) of a dendritic spine and the injected current (of the order of tens of pA) in the spine neck, triggered by a synaptic current.

We apply systematically the electro-diffusion approach to extract the capacitance *C* and the conductance *G* of several spines (SI Fig. S3-5). Using an optimization procedure, we explore the parameter space for computing *C* and *G* (SI Fig. S3-4). We minimize the error between the solution of the electro-diffusion equation and the voltage output of the dendrite during a small time interval at the beginning of the response (SI Fig. S4). The resistance is computed by averaging along the time response over the voltage. We use for the estimator the expression 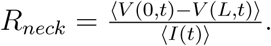 In all cases, we find a good agreement (SI Fig. S5) between the measured and computed voltage drop across the spine neck, where we estimated the current injected in the neck from the head of several spines.

In summary, the average resistance is 〈*R*〉 = 99.2±34.5MΩ (see table 1). To conclude, the electro-diffusion model allows computing the injected current in the spine neck from the head. We also reported here a large variability in the spine resistance, while the capacitance is negligible.

**Table 1:** Parameters extracted from modeling

| Spine | Neck Length L<br>($\mu\text{m}$) | Head radius<br>$r_{head}$ ($\mu\text{m}$) | Intrinsic<br>Capa. C(pF) | Intrinsic<br>Res. $1/G$ ($M\Omega$) | Effective neck<br>Res. $R_{neck}$ ($M\Omega$) |
| --- | --- | --- | --- | --- | --- |
| **1** | 0.58 | 1.18 | $<0.5$ | 128.5 | 73.3 |
| **2** | 0.696 | 0.79 | $<0.5$ | 163.4 | 57.1 |
| **3** | 0.812 | 0.67 | 18 | 212 | 99.15 |
| **4** | 1.044 | 0.75 | 10 | 252 | 132.6 |
| **5** | 1.16 | 0.60 | $<0.5$ | 261 | 134.1 |

### Voltage transduction in a spine and predictions of electro-diffusion

To analyze how a dendritic spine shapes its voltage response to a synaptic input, we simulate the three dimensional PNP equations (see Methods) in a ball of radius *R_head_* and a spine-like geometry. We computed the distribution of the electrical potential for short spine necks, where the head contains two narrow openings: one of radius *r_o_* = 100nm representing the junction with the neck, and the other of radius *r_i_* = 10*nm* that receives the steady current *I_stim_* of positive charges (Fig. 3B). We computed the distribution of positive *c_p_*(*x*) and negative *c_m_*(*x*) charge concentrations as well as the voltage *ϕ*(*x*) when the potential *ϕ* which is grounded to 0 volts at the end of the spine neck, representing the voltage difference induced by the injected current *I_stim_*. We find the distribution of the voltage along the x-axis (blue) Fig. 3B-C when *I_stim_* = 150*pA* is injected in the spherical geometry as shown in Fig. 3B-E: there are two narrow layers due to the small entry and exit, but the injected current induces a 15 mV drop that can propagate to hundreds of nanometers only inside the spine head. Outside these layers, the voltage is quite uniform, leading to a reduced field convection 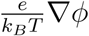 (eq. 8), demonstrating that diffusion is dominant inside the spine head. The regions of large convection are of small sizes (≈ 100*nm* at the limit of actual resolution (≈ 0.116μm/pixel). At this stage, we demonstrated numerically using PNP equations that the voltage drop in the spine head is negligible (less than a quarter of mV), in contrast with the classical cable theory (Fig. S7), which suggests that the motion of ions is driven by the voltage gradient.

**Figure 3:**
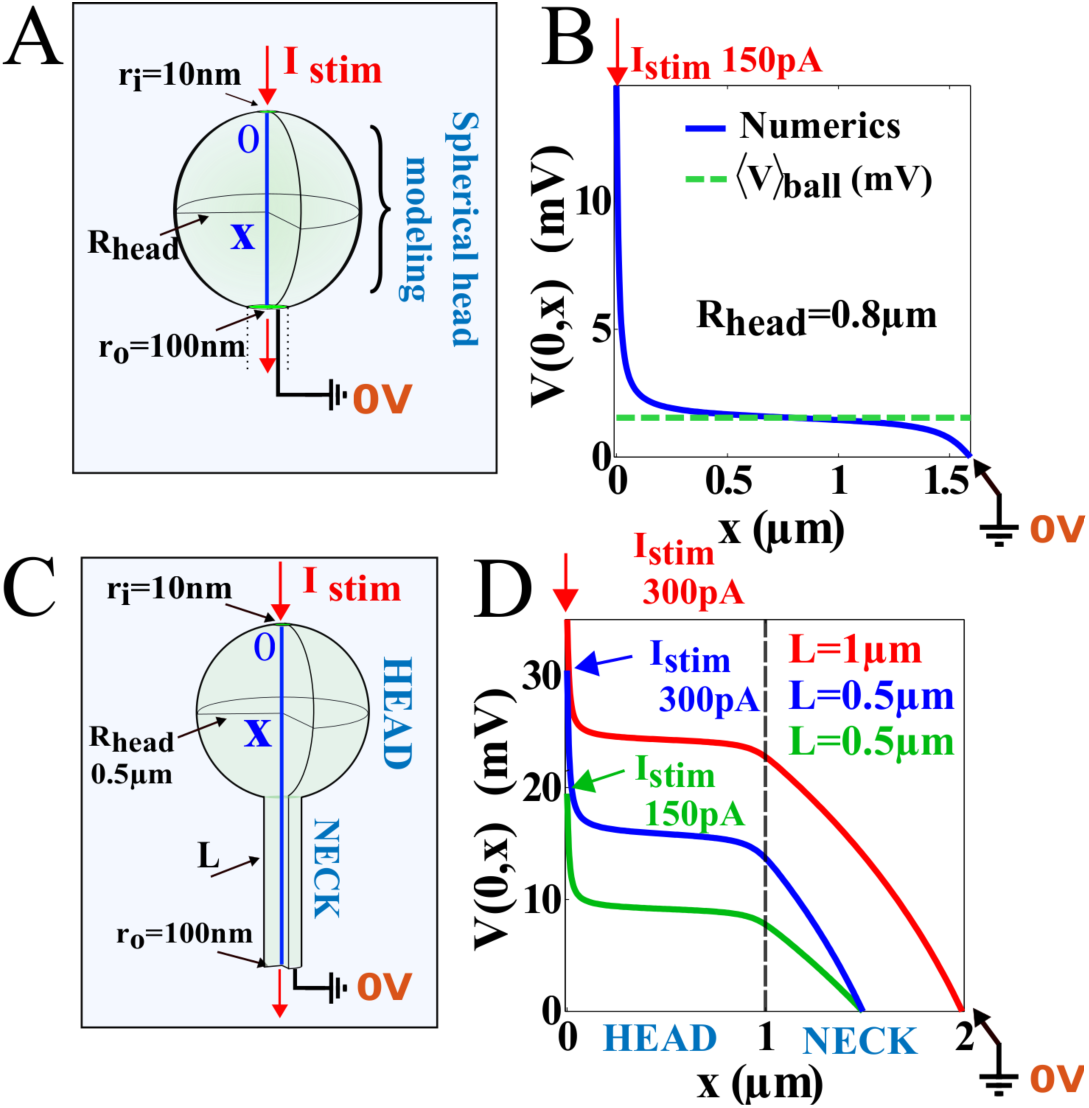
Modeling electro-diffusion in a spine. **A.** Schematic representation of a spine head where a current is injected in a 3D spherical cavity of radius *R_head_* = 0.5*μm*. **B.** Voltage profile (blue) along the x-axis computed from PNP equations, compared to the potential averaged over the entire head (dashed green line) when the injected current is *I_stim_* = 150*pA*. The south pole is grounded at 0 volts. **C.** Schematic representation of a 3D-spine geometry composed by a spherical head of radius *R_head_* = 0.5*μm* and a neck of length *L* = 1*μm*. The head has two narrow openings, one of radius 10*nm* at which the steady current *I_stim_* is injected and a second one of 100*nm* at the junction with the neck. **D.** Potential drop along the x-axis computed from the top of the head to the bottom of the spine. We compare the voltage drop between a spine where *L* = 1*μm* and *I_stim_* = 300*pA* (red) with *L* = 0.5*μm* and *I_stim_* = 150*pA* (blue).

To conclude, inside a spherical domain, diffusion is the dominant driving forces and the potential drop is reduced significantly. We observe that most of the voltage drop is carried by the spine neck (Fig. 3D-E). Interestingly, it is not equivalent to decrease the neck length to compensate for a decrease in the injected current (see result with an injected current of 150 versus 300 pA), suggesting that changing the synaptic weight by adding or removing receptors or modifying spine neck length have different consequences on the spine voltage. At this stage, electro-diffusion theory predicts that the voltage in the spine head *V_head_*(*t*) is spatially homogeneous, supporting the approximation of eq. 1, except near the post-synaptic density or at the entrance of the spine neck. The spine head resistance is thus negligible since the potential drop occurs just at the end of the neck and thus *R_spine_* ≈ *R_neck_*. However, we predicted here that near the post-synaptic density where the current is injected, there should be a nanodomain where the voltage should drop significantly.

### Spine geometry determines the I-V relation

To study the influence of the geometrical parameters on the electrical property of a spine, we first estimated the effect of the spine head radius *R_head_* for five spines (Fig. S5). By measuring their projected area *S_head_* from the two-photon images, we use the relation 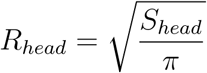 (Fig. 4A) to extract the equivalent radius. We then use the PNP model associated to the short spine with no neck (Fig. 3B) to estimate the average voltage difference (for a current of 100 pA) between the north and the south pole of a spine head 〈*V*〉*_ball_*. We find that the mean voltage varies in a range of 1.5 − 1.6*mV*, when the radius of the head varies in the range 0.3 − 1.5*μm*. This result shows that the head radius had little influence on the mean voltage.

**Figure 4:**
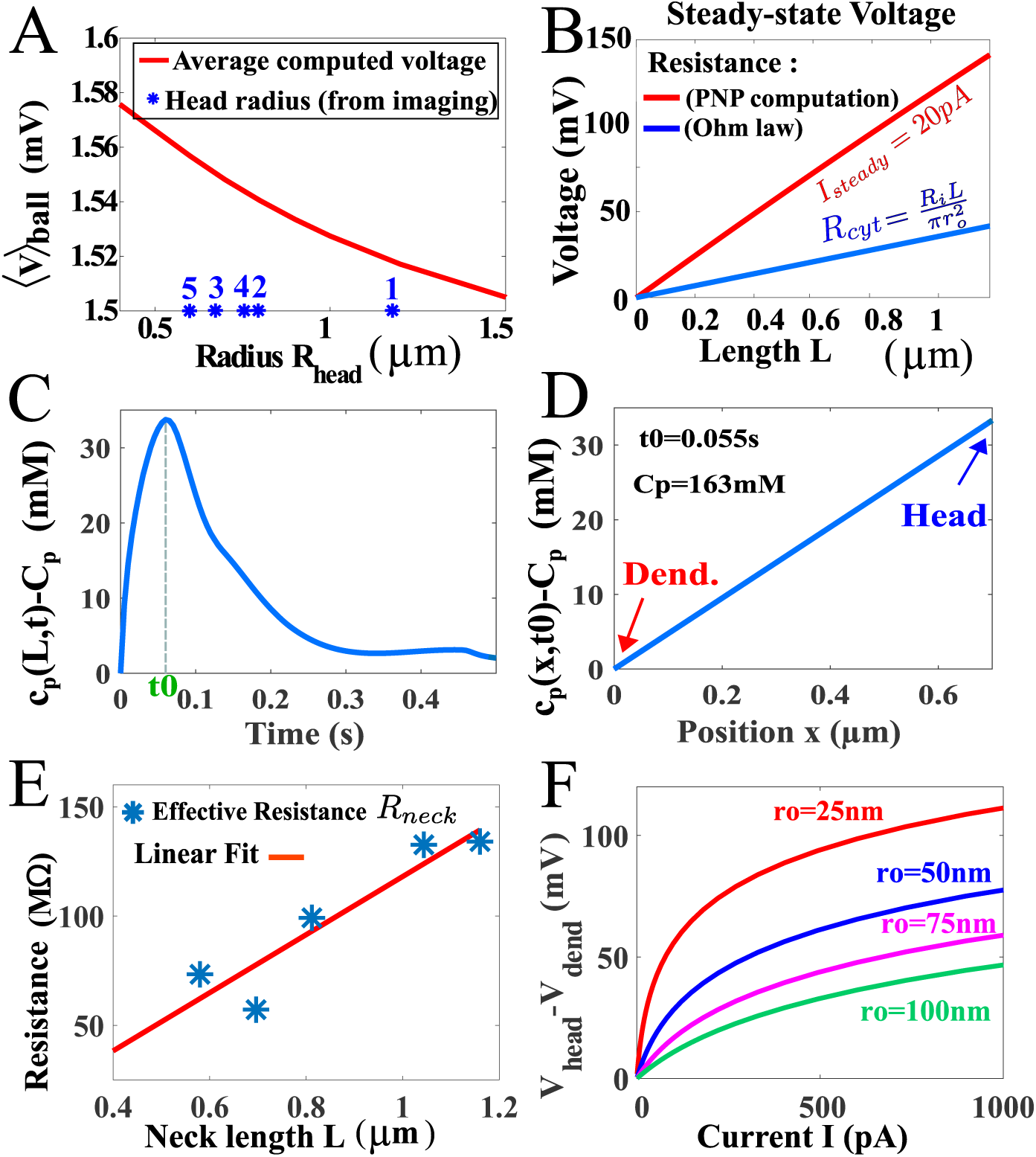
Electrical properties of dendritic spines. **A.** Averaged voltage computed in a spine head using PNP (see fig. 3B): for various head radius, the voltage is almost constant. We position (stars) the estimated radii of different spines head based of their surface, determined from one photon images expressing ArcLight. **B.** Resistance computed at steady state voltage and length of a cylindrical cytosol: we simulate PNP (see Methods) on a wire by injecting a current of 20*pA* that we compare with the classical expression for a resistance [16] with *R_i_* = 109Ω*cm*. **C.** PNP simulation showing the concentration of positive charges at the end of the neck for the injected current described in Fig. 2D. The response peaks at *t*_0_ = 0.055*s*. **D.** Distribution of charges at the peak, computed from PNP, showing a large concentration difference of 33 mM (the total concentration 163 mM). **E.** Estimated spine neck resistance (blue stars) computed as the ratio of the voltage to the current averaged over the time responses *R_neck_* = 〈*V*(0,*t*)〉/〈*I*(*t*)〉 for 5 different spines, revealing how the spine resistance depends on the neck length. **F.** Predicted I-V relation in a dendritic spine for different spine neck radius.

We then investigated the role of the spine neck length (Fig. 4B) when a current of 20*pA* was injected. We compare Ohm’s law with the solution of the PNP equation (ratio voltage/current) in a segment (see Methods). For a resistivity of *R_i_* = 109Ω*cm* (the other parameters are presented in table 1), the difference between the cable and PNP models can be significant as shown in Fig. S7. Furthermore, electrodiffusion in a segment predicts that for the mean current input extracted from data, there is a significant change in the local concentration of positive charges along the segment of length *L* = 0.7*μm* at the time-to-peak (*t*_0_ = 55*ms*). Indeed, the difference in concentration is 33*mM* (the imposed concentration on the other end is 163 mM), leading to a concentration gradient between the dendrite and the spine head (Fig. 4C-D).

Finally, we estimated how the spine neck resistance *R_neck_* depends on the neck length and width, usually unaccessible using classical microscopy approaches: we find both theoretically and experimentally that the resistance increases (blue stars) with the neck length L (Fig. 4E and Fig. 2E). Note that the size of the head is not correlated with the resistance (table 1). Finally, using the electrodiffusion theory and the spine parameter *R_head_* = 0.5*μm*, *L* = 1*μm*, we estimated the I-V relation for various neck radius, showing a saturation for large current (Fig.4F). These curves show that the neck radius is one of the most critical parameter in defining the conversion of current into voltage.

In summary, we used the electro-diffusion theory and the Arclight data to characterize the electrical properties of a dendritic spine. With respect to a synaptic input, a spine can be electrically characterized as a diode device (Fig. 5AB) with a finite resistance (for a small current), saturating for large currents (Fig. 5C). The voltage difference varies from few to tens of mV. However, from the perspective of a Back Propagation Action potential, the equivalent circuit of a spine is a diode with zero resistance (no leak current) Fig. 5D.

**Figure 5:**
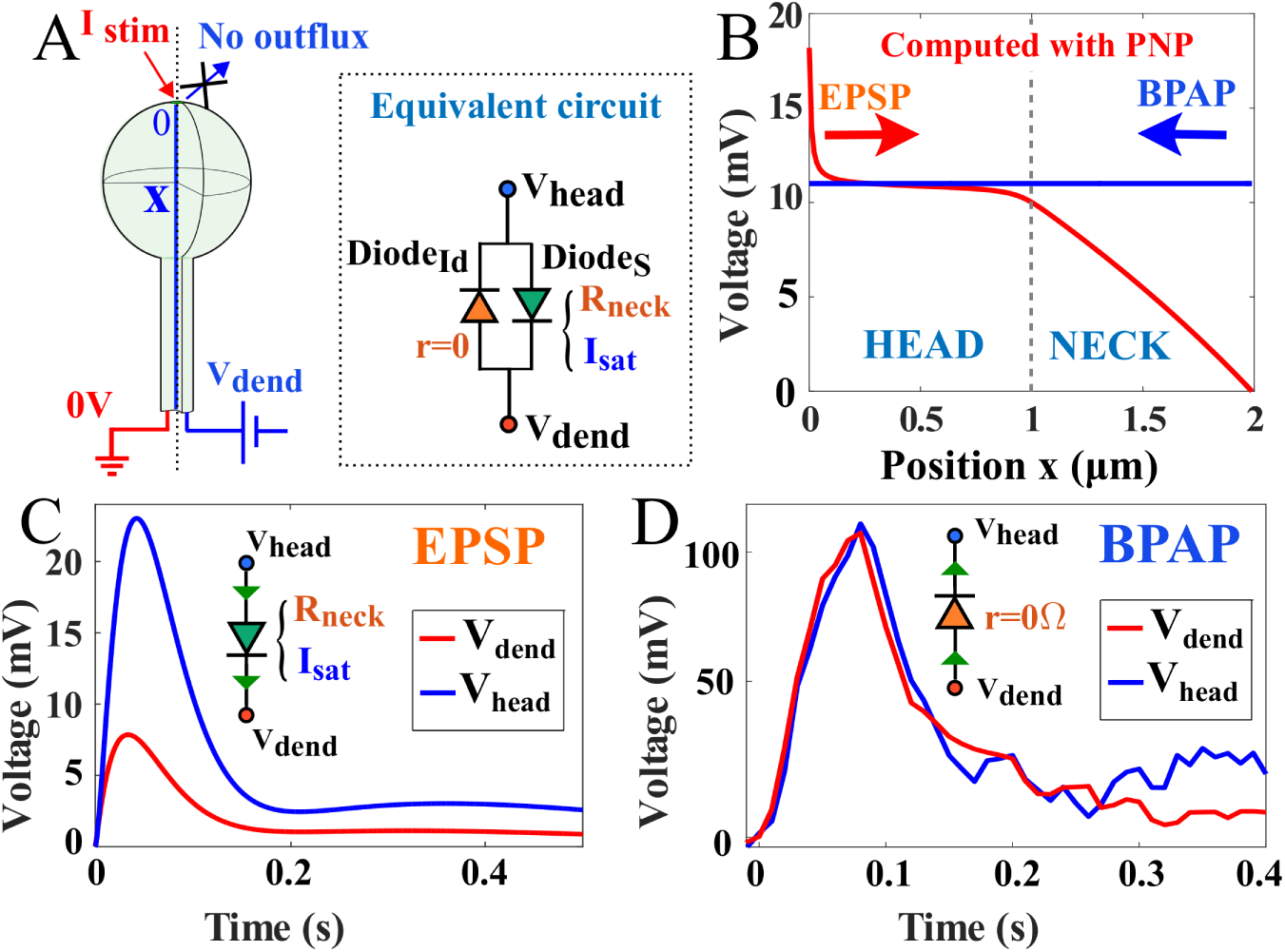
Summary of dendritic spine electrical responses. **A.** Schematic representation of a dendritic spine and equivalent diode circuit where *r* = 0 means that the ideal diode has no resistance. The characteristic of the Shockley diode is *I*_*sat*_, while for the ideal diode, there is no resistance. **B.** Electrical response of a spine (Length and radius of the head are *L* = *R* = 1*μm*) and the radius of the neck *a* = 0.1*μm*) following a synaptic input (*I* = 100*pA*) and a BP action potential (BPAP), where the value depend on the voltage in the dendrite, but it is constant in the spine. **C.** Modulation of the voltage between the spine head and the dendrite: the voltage attenuation can be modeled as a diode to account for the saturation behavior (Fig. 3F). **D.** Response to a BPAP showing no voltage change in the spine.

## Discussion

We developed here a computational approach based on the electro-diffusion theory to estimate the electrical properties of dendritic spines. After we deconvolved the Arclight fluorescent signal, we applied the electro-diffusion theory to estimate the resistance and the capacitance from in vivo hippocampal neuron data. Our approach contrasts with classical estimation of the resistance of the spine neck and the electric properties of dendritic spines in general, that have been extracted in the context of the electrical circuit approximation, cable theory and even diffusion approximation [16, 6]. We found here that the electro-diffusion coupling is the main driving force for the ionic current in the spine neck (Fig. 3), while the diffusion approximation is sufficient to describe the motion of ions inside the head. Indeed, we found that the electric field is negligible in the head, except very close to the entrance of the synaptic input and at the exit with the neck. Electro-diffusion theory reveals that the spine head geometry imposes that the voltage is almost constant in the head, while the neck is responsible for most of the voltage drop. This is in contrast with the predictions of the cable theory or previous approximations of electro-diffusion [16, 4], based on electroneutrality and no gradient of charges. We also demonstrated here that the ion conduction is mostly driven by diffusion in the spine head, suggesting that the head resistance is negligible compared to the neck.

It remains difficult to study the exact local balance of positive by negative charges, because in transient regimes or at equilibrium, positive charges are all the time in excess. Possibly the sum of negatively ionic charges plus the negative charges located on immobile proteins can balance positive charges at a tens to hundreds of nanometers. Long-range electro-diffusion effects have already been described for directing the current flow in the synaptic cleft into the post-synaptic terminal [14, 29], showing in a different context that electro-diffusion drives ionic flows and the voltage in neuronal microdomains.

### Time deconvolution of the Arclight fluorescent signal

Traditional tools to study neuronal voltage are based on recording electrodes. We showed here how the Arclight signal can be deconvolved in small and large microdomains, so that we can now access the voltage dynamics and electrical properties from live cell imaging. Genetically encoded activity sensors combined with novel microscopies are now classically used [30] to record and manipulate the activity of neural circuits. We show here how the fluctuation contained in the fluorescent can be filtered and the voltage time course is recovered from the empirical kernel *K*(*t*) (Methods). This approach can be apply to any encoded activity sensors expressed in neurons and only requires to compare the electrophysiological recording with the florescence in the soma. The present deconvolution could also be used to recover the electrical activity from slow calcium indicators [30].

### Influence of the neck radius on the spine resistance

The spine neck radius cannot be spatially resolved, so any geometrical fluctuation is likely to result in a drastic change in the resistance. For diffusion alone, the rate of extrusion in first approximation is 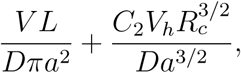 where *D* is the diffusion coefficient, *V* is the volume of the spine, *L* the length of the neck, *R_c_* the radius of curvature at the base of the neck-head junction, *V_h_* the volume of the head and *C*_2_ a constants and *a* is the radius of the neck [31]. This expression shows that a small change in the radius *a* (dividing by two for example) leads to a significant change of at least 4 for the diffusion time scale. We addressed the radius neck uncertainty here in the context of electro-diffusion by computing the neck resistance for different radii (Fig. 3E and S6).

Spine intrinsic electrical characteristics are revealed by the impedance which is the ratio of the voltage to the injected current. For example, for a steady state current of *I* = 50*pA*, the Ohmic resistance of a spine of radius 100*nm* (resp. 50*nm*) is 〈*R_neck_*〉 = 120MΩ, (resp. 〈*R_neck_*〉 = 350MΩ). Interestingly, the spine resistance of a dendritic spine is inversely proportional to the radius of the neck *r*_0_, and not by the square 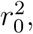 as classically described for electrical devices (SI Fig. S6)[6]. This result shows that the neck size has a key effect in modulating the spine electrical resistance. Another prediction of the present theory is that a synaptic current injected in a spine head should be of the order of 100 pA (as suggested in fig. 3E). Obtaining the shortest diameter of a spine neck along its length is certainly a key factor that could drastically affect its resistance. Indeed, the critical geometrical parameter is the minimal shortest constriction along the neck [31], that could further be influenced by the internal endoplasmic reticulum [21].

### The geometry of dendritic spines modulate the voltage changes independently of the input current

Dendritic spines are involved in modulating two-and three dimensional receptor trafficking [32, 33, 34], molecular post-synaptic density composition, calcium diffusion [5, 7], synaptic transmission and plasticity. We have shown here using the electro-diffusion framework that the voltage in dendritic spines can also be controlled by changing the neck length geometry. This modulation obtained by changing the geometry is complementary to the possible changes in the number of receptors resulting in a long-term modification of the synaptic current, reflecting synaptic plasticity.

Changing the spine neck length can thus regulates the local dendritic voltage, that contributes to the genesis of an action potential. We further confirm previous experimental findings [17, 18], showing that the synaptic amplitude is inversely correlated with the neck length, but we found here a much stronger effect compared to previously evaluated [22]. However, in agreement with [22], we do not need to use any additional active channels in the electro-diffusion model to account for the voltage in the spine, suggesting that they might not play a predominant role.

To conclude, voltage changes in dendrites can now be detected at the nanometer scale and the electro-diffusion theory allows interpreting these data and predicts a nonlinear current-voltage relation imposed by the specific geometry of dendritic spines. While the spine geometry controls voltage, the synaptic current is set by the number of receptors [32, 33, 34]. These two mechanisms are independent and they are both involved in controlling the synaptic response. It would certainly be interesting to study how changes in one affects the other.

## Acknowledgement

DH thanks the hospitality of Department of Applied Mathematics and Theoretical Physics (DAMTP), Churchill College CB3 0DS, this work is supported by a Marie-Curie and a Simons fellowship Award. This research is supported by a DEQ20160334882 Equipes FRM 2016 grant. RY and TK are supported by the NIMH (R01MH101218, R01-MH100561). This material is based upon work supported by, or in part by, the U. S. Army Research Laboratory and the U. S. Army Research Office under contract number W911NF-12-1-0594 (MURI).

## Methods

### Arclight signal

We briefly described here the experimental data we have used for our electrodiffusion theory and time deconvolution. There are fully described in [24]. The protein-based voltage indicator ArcLight is injected in primary cultured hippocampal neurons. ArcLight expressing dissociated hippocampal culture neurons in DIV 12-16 were recorded in artificial cerebrospinal fluid (ACSF) containing ions of various concentration. Two-photon glutamate uncaging was done with a custom-made two-photon laser scanning microscope. In glutamate uncaging, the location of stimulation was selected with 1-2 *μm* distance from dendritic spines, not closer than 1 *μm*. The whole-cell patch clamp and the glutamate uncaging were performing while doing the wide-field one photon imaging of ArcLight fluorescence. Finally, we used the voltage deconvolved from the fluorescence signal, based on a two-state model of voltage dependent ArcLight fluorescence described in [24].

### Deconvolution Kernel

To recover the intrinsic voltage dynamics *h*(*t*) from the slow Arclight signal *G*(*t*), we compare the electrophysiological patch-clamp recording in the soma with the ArcLight fluorescence extracted from the somatic region delimited in the image (Figure 1A). This comparison is at the basis of the deconvolution method of the causal fluorescent signal. Indeed, the slow Arclight reporter convolves the fast electrical voltage signal, modeled by a kernel function *K*(*t*) with the intrinsic dye dynamics, leading to a slow fluorescent response. The kernel *K*(*t*) describes the time delay of the fluorescence activation compared to the voltage dynamics. We model the kernel by the function

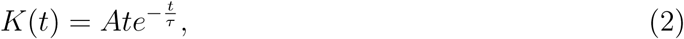

where the value of the parameters *A* and *τ* are obtained by comparing the Arclight response in the soma with the convolution of the electrophysiological recordings (Fig. 2). Indeed, for a voltage signal *h*, the Arclight signal *G*(*t*) is expressed by the convolution product

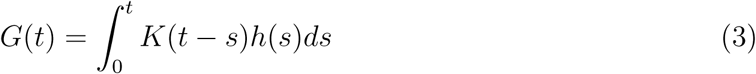

To recovered *h* from the Arclight signal *G*, we first calibrated the kernel so that the Arclight signal peaks exactly at the one monitored by the electrophysiological signal (Figure 1B) and we obtain *τ* = .05*s*. The other parameter *A* is a scaling that will be adjusted for each experimental data. We denote the normalized kernel by 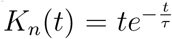 (plotted in SI Fig. 1).

### Noise filtering and approximation

In small dendrite and dendritic spine regions, the Arclight data contains a large noise that should be removed. For that purpose, we use a Savitzky-Golay filter [36], to increase the signal-to-noise ratio. The detail of that procedure is explained in the SI. Once the noise is removed, we define a new step which consists in approximating the signal using a family of analytical functions

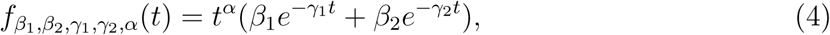

where the parameters *β*_1_, *β*_2_, *γ*_1_, *γ*_2_, *α* are obtained by a best approximation (see SI Fig. S2).

### Microdomain Arclight deconvolution

In the final step, we shall use the deconvolution procedure to compute the voltage. Indeed, once the kernel *K*(*t*) is determined (see above) from the somatic signal and after the step of noise filtering, we shall retrieve the voltage dynamics from dendrites and dendritic spines, where direct electrophysiological recordings are not possible. Using the analytical approximation *G*(*t*) = *f*_*β*_1_,*β*_2_,*γ*_1_,*γ*_2_,*α*_(*t*) of the Arclight fluorescent response (relation 4) described in the previous subsection, we shall now compute the voltage *h* using *K*(*t*) (eq.2) by inverting equation 3 using the Laplace’s transform:

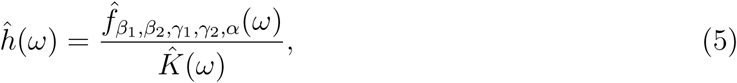

where the Laplace’s transform of the kernel is 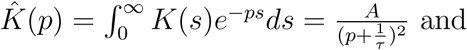

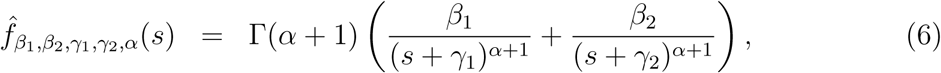

where Г is the Gamma function [38]. The final expression for the voltage is derived in the SI and is given by

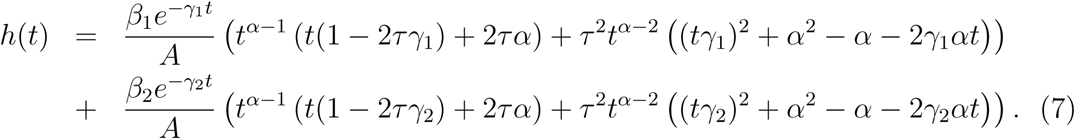

Note that in practice the value of the parameter *A* is calibrated so that the maximum amplitude of the voltage before and after the deconvolution are identical. We applied this procedure to recover the voltage *h* in dendritic spines. This procedure is the time deconvolution of the voltage dynamics from the Arclight fluorescent signal.

### Electro-diffusion model in the spine neck

The Poisson-Nernst-Planck (PNP) equations express the coupling between the ionic flow and the voltage (Poisson equation). We present here the one-dimensional version of these equations. They reduce for the voltage *V* and the concentration of positive *c_p_*(***x***) and negative *c_m_*(***x***) to

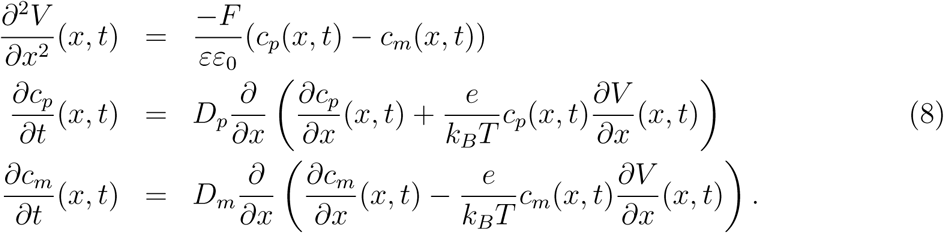

Where *D_p_*, *D_m_* are diffusion coefficients, *e* the electronic charge, the valencies for each specie is *z* = ±1 and *k_B_T* is the thermal energy.

Equations 8-8-9 are used to compute the voltage drop when a current *I*(*t*) is injected at the tip of the spine neck. During the simulations, the ionic concentrations in the dendrite (ionic reservoir) are the boundary conditions fixed at the values *C_p_* and *C_m_* (see table 2). We recall that the electrical potential is defined to an additive constant. The initial and boundary conditions are

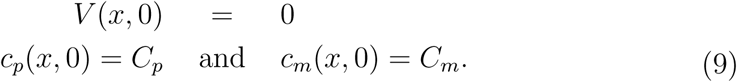

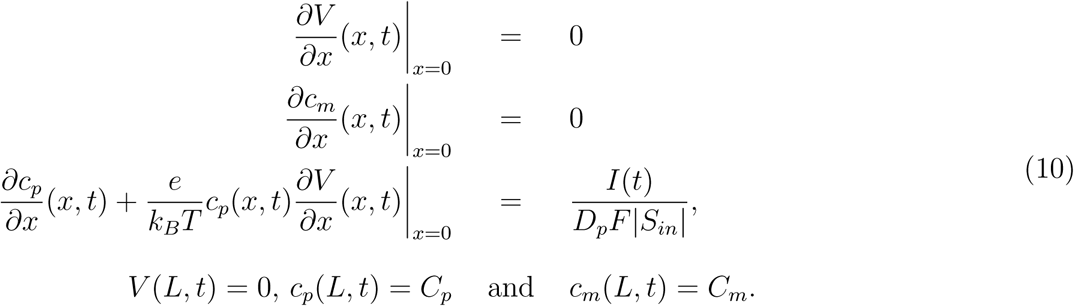

**Table 2:** Biophysical an geometrical parameters.

| Parameter | Description | Value |
| --- | --- | --- |
| $z$ | Valence of ions | 1 |
| $D$ | Diffusion coefficient | $200\mu\text{m}^2/\text{s}$ [37] |
| $D_p$ | Diff. coeff. for + charges | $D$ |
| $D_m$ | Diff. coeff. for – charges | $D$ |
| $C_p$ | + charge concentration | $167\text{mol}/\text{m}^3$ [28] |
| $C_m$ | – charge concentration | $167\text{mol}/\text{m}^3$ [28] |
| $\Omega$ | Spine head | $\Omega$ (volume $\vert \Omega\vert \approx 1\text{fL}$) [5] |
| $a$ | Spine neck radius | (typical) $0.1\mu\text{m}$[35] |
| $L$ | Spine neck length | (typical) $1\mu\text{m}$ |
| $T$ | Temperature | 293.15K |
| $E$ | Energy | $kT = 2.58 \times 10^{-2}\text{eV}$ |
| $e$ | Electron charge | $1.6 \times 10^{-19}\text{C}$ |
| $\varepsilon$ | Dielectric constant | $\varepsilon = 80$ |
| $\varepsilon_0$ | Abs. Dielectric constant | $8.8 \cdot 10^{-12} \text{ F/m}$ |
| $k$ | Boltzmann constant | $1.38 \cdot 10^{-23} \text{ J/K}$ |
| $F$ | Faraday constant | $96485 \text{ As/mol}$ |

In summary, eqs. 8-9-10 describe the ionic response of an input current *I*(*t*) inside a thin cylinder reduced to a one dimensional segment. We simulate these equations using Comsol to determine the voltage drop (Fig. 2).

### 3 dimensional PNP-equations in a Ball and a dendritic spine shape

We present now the steady-state PNP equations, that describe the concentration of positive *c_p_*(***x***) and negative *c_m_*(***x***) charge concentrations and the voltage *ϕ*(***x***) inside a three dimensional bounded domain that we use in Figure 3. The equations are given by

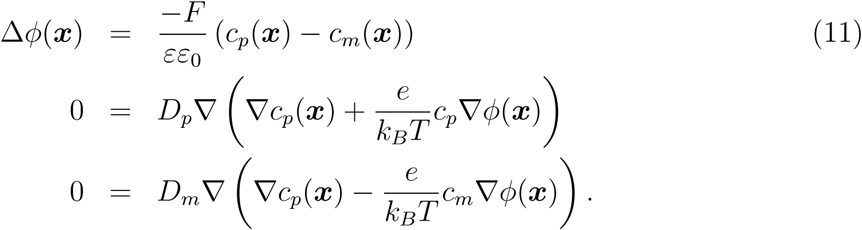

The boundary is decomposed into three subdomains: the current is injected into *∂*Ω_*i*_. Charges can exit in *∂*Ω_*o*_ and the impermeable membrane is represented by *∂*Ω_*r*_. The boundary conditions are

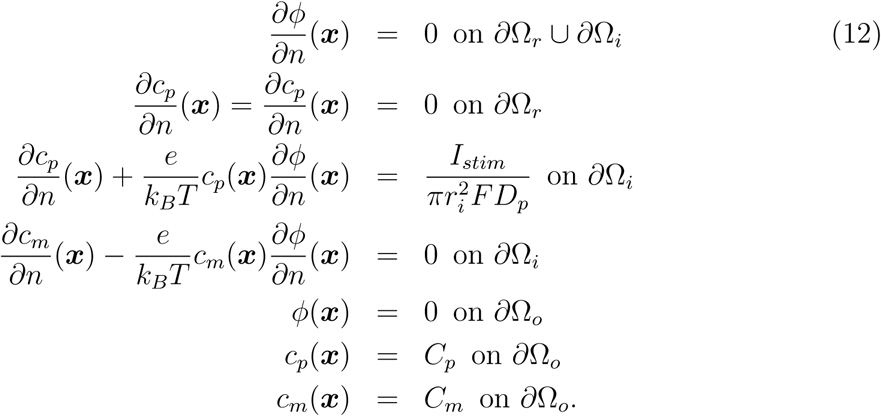

In that model, only positive charges can enter the spine domain. We use the Comsol platform to solve numerically equations 11 presented in fig. 3.

## References

[1] Jack, J.J.B., Noble, D., and Tsien, R.W. Electric current flow in excitable cells. London, Oxford University Press (1975).

[2] Holmes, W.R., Segev, I., Rall, W., Interpretation of time constant and electrotonic length estimates in multicylinder or branched neuronal structures. J Neurophysiol. 68, 1401–20 (1992).

[3] Rall, W. Cable Theory for Dendritic Neurons. In Methods in Neuronal Modeling: from Synapses to Networks, C. Koch, and I. Segev, eds. Cambridge, Mass., MIT Pres, pp. 9–63 (1989).

[4] Qian, N., Sejnowski, T.J. An electro-diffusion model for computing membrane potentials and ionic concentrations in branching dendrites, spines and axons. Biol Cybern 62, 1–15 (1989).

[5] Yuste, R., Dendritic spine, The MIT Press, (2010).

[6] Svoboda, K, Denk, W, Kleinfeld, D, Tank, D.W., In vivo dendritic calcium dynamics in neocortical pyramidal neurons. Nature 385, 161–5 (1997).

[7] Korkotian, E, Holcman, D, Segal M., Dynamic regulation of spine-dendrite coupling in cultured hippocampal neurons. Eur J Neurosci. 20, 2649–2663 (2004).

[8] Bloodgood, B.L., Sabatini, B.L. Neuronal activity regulates diffusion across the neck of dendritic spines. Science 310, 866–9 (2005).

[9] Tonnesen, J., Katona, G., Rozsa, B. Nagerl, U.V. Spine neck plasticity regulates compartmentalization of synapses. Nature neuroscience 17, 678–85 (2014).

[10] Bezanilla, F. How membrane proteins sense voltage. Nat Rev Mol Cell Biol. 9, 323–32 (2008).

[11] Eisenberg RS., Look at biological systems through an engineer’s eyes. Nature. 2007 May 24;447(7143):376.

[12] Eisenberg, R.S. From structure to function in open ionic channels. J Membr Biol 171, 1–24 (1999).

[13] Eisenberg, R.S., M.M. Klosek, and Z. Schuss, (1995). “Diffusion as a chemical reaction: Stochastic trajectories between fixed concentrations.” J. Chem. Phys., 102 (4), pp.1767–1780.

[14] Sylantyev, S., Savtchenko, L.P., Ermolyuk, Y., Michaluk, P. Rusakov, D.A. Spike-driven glutamate electrodiffusion triggers synaptic potentiation via a homer-dependent mGluR-NMDAR link. Neuron 77, 528–41 (2013).

[15] Savtchenko, L.P., Kulahin, N., Korogod, S.M., Rusakov, D.A. Electric fields of synaptic currents could influence diffusion of charged neurotransmitter molecules. Synapse 51, 270–278 (2004).

[16] Koch, C. Biophysics of Computation: Information Processing in Single Neurons. Oxford University Press, USA; New Ed edition (2004).

[17] Araya, R., Jiang, J., Eisenthal, K.B., Yuste, R. The spine neck filters membrane potentials. Proc. Natl. Acad. Sci. USA 103, 17961–17966 (2006).

[18] Araya, R., Vogels, T.P., Yuste, R. Activity-dependent dendritic spine neck changes are correlated with synaptic strength. Proc. Natl. Acad. Sci. USA 111, 2895–904 (2014).

[19] Hochbaum, D.R., Zhao, Y., Farhi, S.L., Klapoetke, N., Werley, C.A., Kapoor, V., Zou, P., Kralj, J.M., Maclaurin, D., Smedemark-Margulies, N., Saulnier, J.L., Boulting, G.L., Straub, C., Cho, Y.K., Melkonian, M., Wong, G.K., Harrison, D.J., Murthy, V.N., Sabatini, B.L., Boyden, E.S., Campbell, R.E., Cohen, A.E., All-optical electrophysiology in mammalian neurons using engineered microbial rhodopsins. Nat Methods. 11, 825–33 (2014).

[20] Harnett, M.T., Makara, J.K., Spruston, N., Kath, W.L., Magee, J.C., Synaptic amplification by dendritic spines enhances input cooperativity. Nature 491, 599–602 (2012).

[21] Holcman, D., Yuste, R., The new nanophysiology: regulation of ionic flow in neuronal subcompartments. Nat Rev Neurosci. 16, 685–92 (2015).

[22] Popovic, M.A., Carnevale, N., Rozsa, B., Zecevic, D. Electrical behaviour of dendritic spines as revealed by voltage imaging. Nat Commun. 6, 8436 (2015).

[23] Acker, C. D., Hoyos, E., Loew, L. M. EPSPs Measured in Proximal Dendritic Spines of Cortical Pyramidal Neurons. eNeuro 3, doi:10.1523/ENEUR0.0050-15.2016 (2016).

[24] T. Kwon, M Sakamoto, D. Peterka and Yuste, R. Voltage compartmentalization by dendritic spines (revision) (2016).

[25] Jackson, J.D. Classical Electrodynamics (3rd ed.). New York: John Wiley & Sons (1998).

[26] Rubinstein, I., Electro-Diffusion of ions. SIAM, Studies in Applied Mathematics (1987).

[27] Andelman, D. Electrostatic Properties of Membranes: The Poisson-Boltzmann Theory, Elsevier Science B.V. Handbook of Biological Physics (1995).

[28] Hille, B. Ionic Channels in Excitable Membranes. 3nd edn. Sunderland, Mass., Sinauer. (2001).

[29] Sylantyev, S., Savtchenko, L.P., Ermolyuk, Y., Michaluk, P. Rusakov Electric fields due to synaptic currents sharpen excitatory transmission. Science 319, 1845–1849 (2008).

[30] Emiliani V, Cohen AE, Deisseroth K, Husser M., All-Optical Interrogation of Neural Circuits. J Neurosci. 2015;35(41):13917–26.

[31] Holcman, D. & Schuss, Z. Control of flux by narrow passages and hidden targets in cellular biology, Phys. Progr. Report 76, 7 (2013).

[32] Huganir, R.L., Nicoll, R.A. AMPARs and synaptic plasticity: the last 25 years. Neuron 80, 704–17 (2013).

[33] Elias, G.M., Nicoll, R.A. Synaptic trafficking of glutamate receptors by MAGUK scaffolding proteins. Trends Cell Biol. 17, 343–52 (2007).

[34] Kessels, H.W., Malinow, R. Synaptic AMPA receptor plasticity and behavior. Neuron 61, 340–50 (2009).

[35] Takasaki, K.T., Ding, J.B., Sabatini, B.L. Live-cell superresolution imaging by pulsed STED two-photon excitation microscopy. Biophys. J. 104, 770–777 (2013).

[36] Savitzky, A., Golay, M. J. E., Smoothing and Differentiation of Data by Simplified Least Squares Procedures. Anal. Chem. 36, 1627–39 (1964).

[37] Chen, K. C. & Nicholson, C. Spatial Buffering of Potassium Ions in Brain Extracellular Space, Biophys. J. 78, 277697 (2000).

[38] Abramowitz, M., & Stegun, I. A. Handbook of Mathematical Functions. Dover Publications Inc.; New edition edition (1965).

